# Yeast-MetaTwin for Systematically Exploring Yeast Metabolism through Retrobiosynthesis and Deep Learning

**DOI:** 10.1101/2024.09.02.610684

**Authors:** Ke Wu, Haohao Liu, Manda Sun, Runze Mao, Yindi Jiang, Eduard J. Kerkhoven, Yu Chen, Jens Nielsen, Feiran Li

## Abstract

Underground metabolism plays a crucial role in understanding enzyme promiscuity, cellular metabolism, and biological evolution, yet experimental exploration of underground metabolism is often sparse. Even though yeast genome-scale metabolic models have been reconstructed and curated for over 20 years, more than 90% of the yeast metabolome is still not covered by these models. To address this gap, we have developed a workflow based on retrobiosynthesis and deep learning methods to comprehensively explore yeast underground metabolism. We integrated the predicted underground network into the yeast consensus genome-scale model, Yeast8, to reconstruct the yeast metabolic twin model, Yeast-MetaTwin, covering 16,244 metabolites (92% of the total yeast metabolome), 2,057 metabolic genes and 59,914 reactions. We revealed that *K*_m_ parameters differ between the known and underground network, identified hub molecules connecting the underground network and pinpointed the underground percentages for yeast metabolic pathways. Moreover, the Yeast-MetaTwin can predict the by-products of chemicals produced in yeast, offering valuable insights to guide metabolic engineering designs.

## Introduction

Underground metabolism refers to biochemical processes that are currently inadequately understood, often metaphorically referred to as the "dark matter" of metabolism^1^. It mainly includes the activities of enzymes that exhibit broad substrate specificity—known as enzyme promiscuity^2^—and the roles of uncharacterized enzymes remain to be discovered. This flexibility of underground metabolism allows enzymes to evolve new functions, helping organisms adapt to different environmental conditions^3^. However, this can also lead to undesired side effects, such as the degradation of natural or heterologous products by unknown enzymes during cell factory engineering^4–6^.

Genome-scale metabolic network models (GEMs) are widely utilized to systematically study cell metabolism^7^. These models describe the gene-protein-reaction associations of metabolic functions of an organism, and have proven instrumental in studying cellular phenotypes, designing cell factories, and analyzing omics data^8, 9^. Thus, significant efforts have been dedicated in the development and curation of the GEMs^10, 11^. However, taking the model organism *Saccharomyces cerevisiae* as an example, despite 20-year refinement, 14,882 metabolites with SMILES reported in the yeast metabolome database (YMDB)^12^ are still missing in the consensus yeast GEM-Yeast8^13^. This indicates that there remains an extensive uncharacterized biochemical space in our current understanding of yeast metabolism, leading to erroneous predictions and potentially flawed applications in biotechnology and biomedicine^14^.

To date, experimental exploration has covered only a small fraction of the underground metabolism^15^, focusing primarily on the promiscuity of individual enzymes^3, 16–18^. Although computational methods such as retrobiosynthesis prediction bring promise for identifying unclear pathways^19–22^, their effectiveness is limited by the vast number of possible combinations, making it difficult to identify successful pathways and their associated enzymes^23, 24^. Consequently, these methods mainly serve as gap-filling tools within metabolic models^25, 26^. Recently, deep learning have significantly improved enzyme function annotation, demonstrating strong performance in predicting parameters, such as *k*_cat_ ^27–29^, *K*_m_ ^28, 30^, EC numbers^31, 32^ and enzyme-substrate pairs^33^. However, integrating these advancements with the study of underground metabolism remains unexplored. This presents an opportunity for developing a systematic and effective workflow to bridge this gap and enhance our understanding of underground metabolic processes, as well as their applications for understanding metabolic mechanisms^34^, biological evolution^2, 35^, and guiding the natural products production^36, 37^.

To this end, we developed a workflow that integrates rule-based retrobiosynthesis reaction prediction with deep learning-based enzyme annotation. Using this approach, we systematically explored yeast underground metabolism by managing the combinatorial explosion of reaction possibilities and annotating enzymes for the predicted reactions. This resulted in the development of Yeast-MetaTwin, a comprehensive yeast metabolic model. Additionally, we predicted kinetic parameters to reveal differences between known and underground metabolism and demonstrated the unique applications of Yeast-MetaTwin in guiding engineering strategies for cell factory design.

## Results

### The pipeline to generate comprehensive yeast metabolic reaction dataset

We first compared the 16,042 metabolites in YMDB with the metabolites in Yeast8 and identified 14,882 metabolites missing from Yeast8. According to their classification via ClassyFire^38^, these missing metabolites comprise 14,310 lipids and 572 non-lipids (Fig. 1a). Among the non-lipid metabolites, the most common classes included organoheterocyclic compounds, benzenoids, organic acids and derivatives (Fig. 1b). To explore the metabolic network connecting these missing reaction links among metabolites, we designed a workflow for systematic mining of underground metabolism (Fig. 1c). Specifically, we first extracted 21,921 known biochemical reaction rules from biochemical reaction database-MetaNetX^39^ and used these rules as templates to expand possible reactions among all yeast metabolites (Supplementary Table 1 and Supplementary Figure 1). This resulted in around 179 million reactions among lipid metabolites and 4 million reactions for non-lipid metabolites. Next, we extracted a connected yeast network with 1,092,946 reactions from this reaction pool by removing dead-end reactions and ensuring that all metabolites involved in these reactions exist in the yeast metabolome dataset. By assuming that the yeast metabolome covers all metabolites in yeast, we can constrain the reaction expansion search to a single step and exclude reactions involving metabolites outside this metabolite dataset. This strategy simplifies the originally complex process of analyzing multiple step combinations—whose complexity grows exponentially—by reducing it to the analysis of single step combinations, which has quadratic complexity. This approach effectively covers nearly all possible combinations. It streamlines the analysis by avoiding the need to account for intermediate metabolites that are not present in yeast and by not having to consider pathway length, which is a common issue with traditional retrobiosynthetic methods.

**Figure 1.**
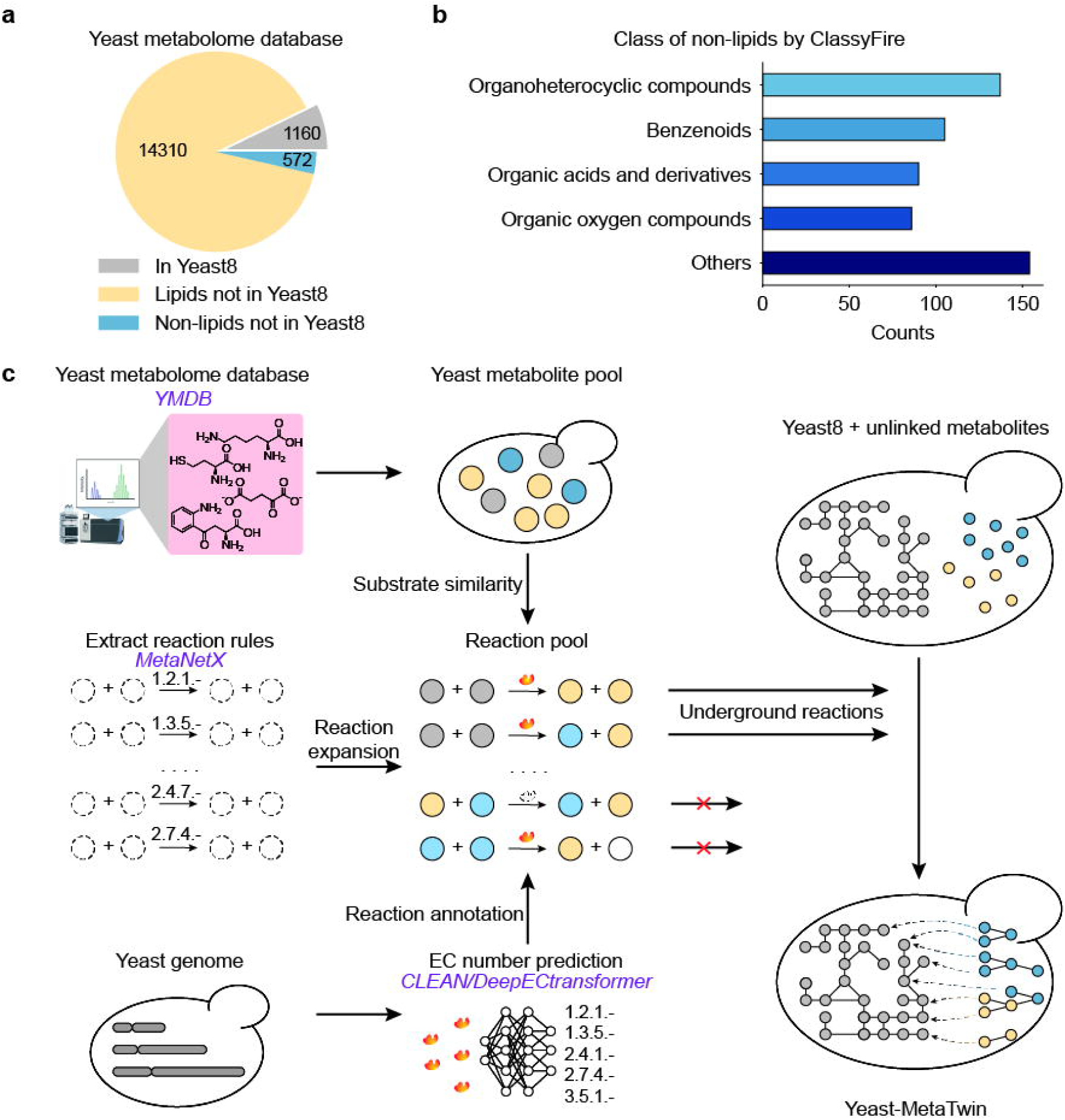
Mining the comprehensive metabolism of *S. cerevisiae*. (a) Comparison of metabolites between Yeast8 and YMDB database. (b) Classification of non-lipid metabolites in YMDB that are not included in Yeast8. (c) Workflow for mining yeast comprehensive metabolism from metabolome data based on retrobiosynthesis and deep learning.

Next, we annotated enzyme capable of catalyzing those predicted reactions. We employed two deep learning-based EC number prediction methods, CLEAN^31^ and DeepECtranformer^32^, to predict EC numbers for all yeast genes. In total, 2,445 genes were annotated with 161 reaction types, categorized by the first three digits of the EC number (e.g., EC X.X.X.-). Then, we mapped these genes to those predicted reactions based on EC number matching (Fig. 1c). Predicted reactions inherited the first three EC number digits from the template reaction rules, thus those reaction types can be used to map the gene to the reactions^40, 41^. Reactions lacking successful gene annotations were excluded, indicating that yeast is not likely to catalyze these reactions, and they should therefore not be included in the model. The remaining gene-annotated and fully connected reaction network with 59,914 reactions, 16,244 metabolites and 2,057 genes (covering 84% of the predicted enzymes) represent the comprehensive yeast metabolic network. The detailed process for reaction prediction and annotation can be found in Supplementary Figure 2.

The effects of parameters on performance were evaluated by calculating the recovery rate, which measures the proportion of known yeast reactions accurately captured in the predicted comprehensive reaction dataset. We found that this recovery rate can be improved by lowering the metabolite similarity cutoff and including more similar metabolites for reaction expansion (Supplementary Figure 3a). However, to enhance computational efficiency during the reaction prediction step and reduce the increased false discovery rate in gene annotation, we set a substrate similarity score cutoff > 0.3 and applied this criterion to the top 50 matched substrates for reaction expansion (Supplementary Figure 3b). With the determined parameters, we found that approximately 78% of the Yeast8 reactions were recovered in our predictions, demonstrating the predictive power of this workflow (Supplementary Figure 3a).

As for the enzyme annotation, CLEAN and DeepECtransformer successfully annotated 86% and 66% of these reactions compared with enzyme annotation in Yeast8, respectively (Supplementary Figure 4a). Considering that the CLEAN method brings a large number of false discovery results (Supplementary Figure 4a-b), we combined the two methods as the final annotation scheme, which effectively reduces the false discovery rate while maintaining the high recovery rate of CLEAN (Supplementary Figure 4a). Furthermore, to verify that the high recovery rate is not resulting from human bias in the Yeast-GEM reconstruction and curation processes^13^, we further compared our predictions with yeast annotations in UniProt database and identified a high recovery rate for reactions (79%) and for enzyme annotations (85%) for *S. cerevisiae* annotation in UniProt (Supplementary Figure 5), demonstrating the robustness of the pipeline in identifying the known metabolic network of yeast metabolism and thus high confidence of the predicted yeast reaction dataset.

### Reconstruction and performance of the comprehensive yeast metabolic twin model: Yeast-MetaTwin

In addition to the 4,131 reactions that already existed in Yeast8 (known reactions), there are 55,783 unique reactions (underground reactions) in the final predicted yeast metabolic network, demonstrating that 93% of the yeast reaction network has not previously been explored through systematic modeling. Among the 55,783 reactions, 3,381 are related to non-lipid metabolites and 52,402 are involving lipids.

First, we expanded Yeast8 with the 55,783 predicted underground reactions to generate the comprehensive yeast metabolic twin model, named Yeast-MetaTwin. This model serves as a twin representation of the yeast metabolism, since it represents the most comprehensive assembly of metabolic conversions that can happen in yeast. Yeast-MetaTwin contains 2,057 genes, 16,244 metabolites, and 59,879 reactions (Fig. 2a).

**Figure 2.**
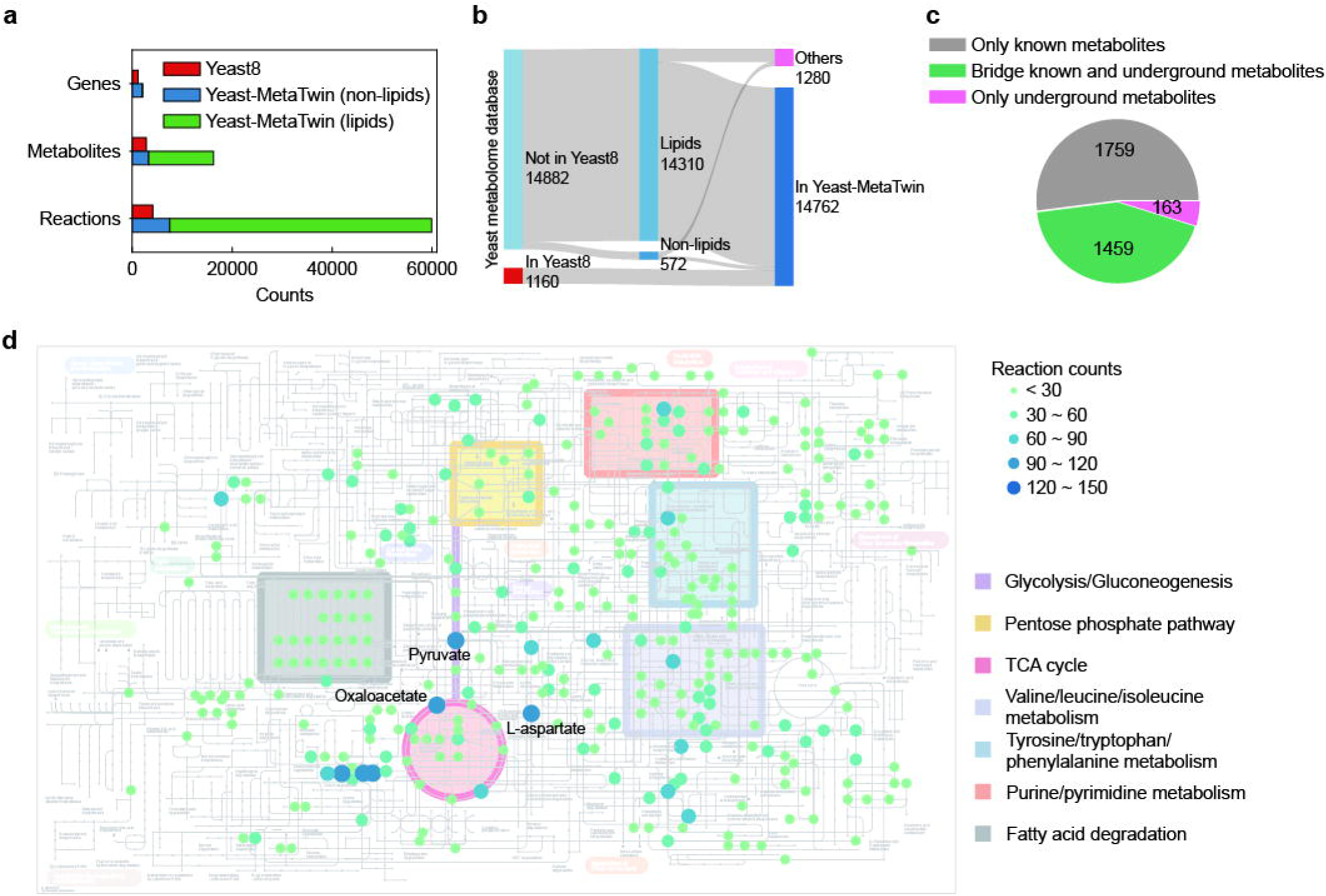
Parameters of the Yeast-MetaTwin and distributions of underground metabolism. (a) Statistical comparison between Yeast8 and Yeast-MetaTwin. (b) Successfully linked metabolites of the yeast metabolite pool in Yeast-MetaTwin (c) Percentage of underground metabolism reactions categorized by metabolite involvement: reactions involving only Yeast8 metabolites, reactions involving only previously unexplored metabolites, and reactions bridging both Yeast8 and previously unexplored metabolites. Lipid-related reactions are excluded from this analysis. (d) Count of reactions associated with each metabolite in underground metabolism (excluding lipids). In the metabolic maps, nodes represent metabolites from Yeast8 involved in underground metabolism, with their color and size corresponding to the number of reactions they participate in.

Then, we validated the predictive capabilities of Yeast-MetaTwin. We simulated the various yeast phenotypes including growth, BIOLOG substrates utilization and gene essentiality. Compared with Yeast8, Yeast-MetaTwin showed improved performance in gene essentiality (Accuracy: 89% for Yeast8, 92% for Yeast-MetaTwin) and growth simulation (Pearson ’s *r*: 0.50 for Yeast8, 0.61 for Yeast-MetaTwin), and similar performance in BIOLOG utilization (Accuracy: 82% for Yeast8, 81% for Yeast-MetaTwin), demonstrating its ability with precise predictions even with significant model expansion (Supplementary Figure 6a-c). Additionally, we examined yeast robustness to gene function loss and its impact on phenotypes. Studies have shown that various bacteria can still synthesize essential amino acids and grow on minimal media through underground metabolic activities, even if their genomes lack the genes typically required for amino acid synthesis^42^. It has been shown that bacteria can still synthesize amino acids unless more than 40% of the genes in the amino acid synthesis pathways are knocked out^43^. To assess the robustness of amino acid synthesis in yeast, we used Yeast8 and Yeast-MetaTwin models to simulate the ability of yeast to retain amino acid synthesis when different proportions of genes in amino acid synthesis pathways were knocked out. According to the Yeast8, yeast lost the ability to synthesize all amino acids when the gene knockout percentage was 0.2 (Supplementary Figure 6d). However, Yeast-MetaTwin showed that yeast would completely lose the ability when the gene knockout percentage reached 0.4 (Supplementary Figure 6d). Considering yeast underground metabolism resulted in a similar gene knockout percentage to what has been experimentally observed in bacteria, where underground metabolism also plays a critical role.

Compared to previous yeast GEMs, and even GEMs for any species, Yeast-MetaTwin has a hugely expanded metabolite coverage. Out of the 14,310 lipid and 572 non-lipid YMDB metabolites that were not present in Yeast8, we successfully filled the knowledge gaps for 13,336 lipid and 266 non-lipid metabolites, together with the GPR associations (Fig. 2b). This effort has substantially increased the exploration rate of the yeast metabolome from 7% to 92%. To manage the computational burden and maintain efficiency, and to prevent the vast number of lipid metabolism-related reactions from overshadowing those of non-lipid metabolism, we present lipid and non-lipid metabolism separately in the following analyses. Among the connected 266 non-lipid metabolites, 215 were successfully annotated with GPRs without introducing any out-of-yeast metabolites; 51 were activated by adding exchange and transport reactions since they are documented as obtained from plants and environment in YMDB (Supplementary Table 2). Among the metabolites that were not successfully connected, four could be synthesized when reactions without GPR annotations were allowed, without introducing any non-yeast metabolites (Supplementary Table 3). Additionally, 267 metabolites were successfully connected by including one or two non-yeast metabolites as co-substrates or co-products. This suggests the presence of potential metabolites yet to be identified in yeast and provides valuable clues for further yeast metabolomics research (Supplementary Table 4). As for the remaining unconnected metabolites, majority of them cannot be matched to any reaction rules in our pipeline, which was also confirmed as unsuccessful by other retrobiosynthesis methods, including deep learning-based methods such as RetroPathRL^40^ and ASKCOS^44^ (Supplementary Table 5). This suggests the presence of unexplored reaction mechanisms or incorrect reporting on the yeast metabolome.

### Hub metabolites are widely spread among the metabolic networks

Among the reactions of yeast underground metabolism, we identified 52.0% as new links among known Yeast8 metabolites, 4.8% in unexplored metabolites (not present in Yeast8) and the remaining bridging the known and the unexplored metabolites (Fig. 2c). This suggests that underground metabolism primarily underlines the known metabolites with previously hidden links. Furthermore, we used iPath3^45^ to visualize the underground metabolism to reveal metabolite frequencies within underground metabolism. Underground activities are prevalent in cellular metabolism (Fig. 2d). The metabolites with more frequent underground activities, termed as “hub compounds”, include nucleotide metabolites, amino acids and central carbon metabolism (such as pyruvate and oxaloacetate) (Fig. 2d). The pattern in the underground metabolism is preserved in the Yeast-MetaTwin model (Supplementary Figure 7). We can see those metabolites matched to more reaction rules are more likely to be a hub compound (Supplementary Figure 8, *r* = 0.41 for compounds with carbon number 3-9, *P* < 0.001). Regarding lipid-related underground metabolism, the main pattern also preserved, but with L-methionine being the most active hub metabolites (Supplementary Figure 9), which is likely due to L-methionine’s prevalent roles in phospholipid synthesis and methylation processes related to lipid metabolism^46^.

### Affinity differs underground from the known Yeast8 metabolism

To explore the kinetic characteristics of Yeast8 (known metabolism) and underground metabolism (predicted new reactions), we utilized published deep learning methods to predict the catalytic rate constants (*k*_cat_) and Michaelis constants (*K*_m_) for the known and underground networks. We employed UniKP-*k*_cat_ ^28^, TurNuP^29^, and DLKcat^27^ for *k*_cat_ prediction; while UniKP-*K* ^28^ and an unnamed machine learning prediction method^30^, that we refer to as “Boost_KM”, for *K*_m_ prediction.

Both DLKcat and UniKP-*k*_cat_ are *k*_cat_ prediction methods based on enzyme-substrate pairs. For DLKcat, the median *k*_cat_ values are 5.71 s⁻¹ in underground metabolism and 5.27 s⁻¹ in known metabolism (Fig. 3a). For UniKP-*k*_cat_, the median *k*_cat_ values are 3.41 s⁻¹ in underground metabolism and 4.08 s⁻¹ in known metabolism (Fig. 3b). Overall, the distribution of *k*_cat_ values for known enzymes is similar to that of the underground network from both prediction methods. TurNuP predicts the *k*_cat_ value of a reaction based on the enzyme sequence and the complete reaction SMILES, rather than the substrate. For the underground metabolic network, the median *k*_cat_ value is 10.71 s^-1^, while for known metabolism it is 10.98 s^-1^, which also indicates the similar *k*_cat_ distribution between the known and underground datasets (Fig. 3c).

**Figure 3.**
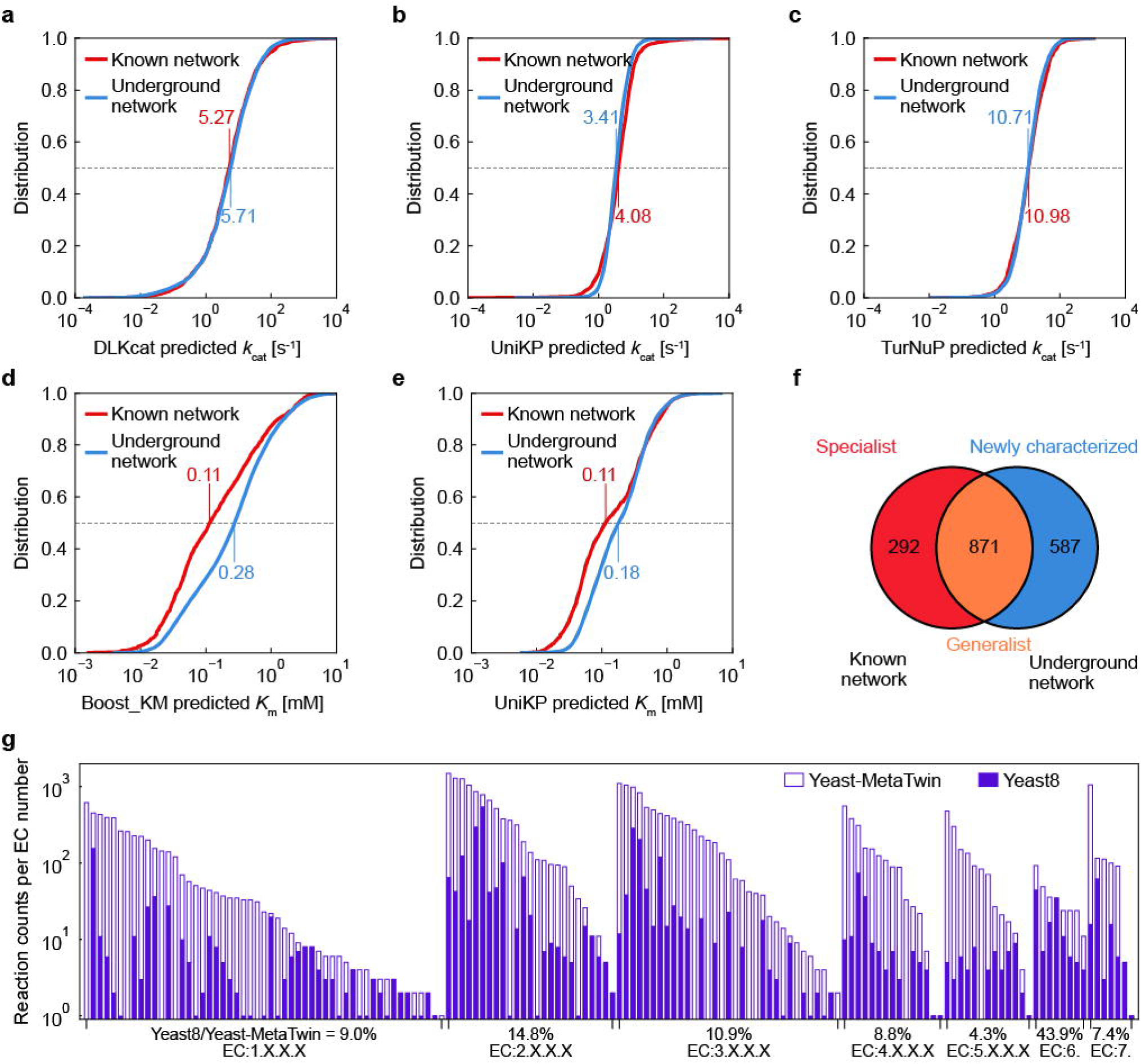
Kinetic characteristics and enzyme classification distributions of known and underground network. Kinetic parameter prediction in Yeast-MetaTwin for both the known network (Yeast8) and underground networks (predicted new reactions): (a) distribution of *k*_cat_ values predicted by DLKcat, (b) distribution of *k*_cat_ values predicted by UniKP-*k*_cat_, (c) distribution of *k*_cat_ values predicted by TurNuP, (d) distribution of *K*_m_ values predicted by Boost_KM, and (e) distribution of *K*_m_ values predicted by UniKP-*K*_m_. (f) Venn diagram illustrating gene sets from the known and underground network in Yeast-MetaTwin. (g) Proportions of explored reactions in different EC number categories.

Both Boost_KM and UniKP-*K*_m_ predict *K*m values based on enzyme-substrate pairs. For these methods, the median *K*m values in the underground and known metabolic networks are 0.28 mM/0.11 mM (Boost_KM) and 0.18 mM/0.11 mM (UniKP-*K*_m_), showing approximately a 2-fold difference in median values (Fig. 3d-e). Overall, the *K*_m_ values in underground metabolism are consistently higher than those in the known metabolic network, indicating that *K*_m_ is a key parameter distinguishing these two metabolic networks. In summary, *K*_m_ values more effectively differentiate between underground and known metabolic networks, and the consistent results obtained from various prediction models confirm the reliability of this conclusion.

To better understand the source of this variation, we divided enzyme involved in underground metabolism into two categories: enzymes with promiscuous activity and previously uncharacterized enzymes. Among the total 2,057 enzymes in the Yeast-MetaTwin, 307 are associated with either transport reactions or pseudo-reactions, which do not contain specific SMILES information and therefore cannot be used for kinetic predictions. For the remaining 1,750 enzymes in the Yeast-MetaTwin, 292 are specialist enzymes from Yeast8, which retain their original functions without participating in any novel underground reactions (Fig. 3f). In contrast, 871 from Yeast8 exhibit enzyme promiscuity, while 587 represent previously uncharacterized enzymes. We analyzed the kinetic parameters across four distinct categories: reactions catalyzed by Yeast8 specialist enzymes (292 enzymes), previously known reactions catalyzed by Yeast8 generalist enzymes (871 enzymes), underground reactions catalyzed by Yeast8 generalist enzymes (871 enzymes), and underground reactions catalyzed by previously uncharacterized enzymes (587 enzymes). We identified that both components of the underground metabolic network—enzyme promiscuity of known enzymes and newly characterized enzymes—exhibited consistent distributions of observation captured in Fig. 3a-e (Supplementary Figure 10a-e). This consistency suggests that both parts contribute to the observed differences in *K*_m_ values between underground and known metabolism. Moreover, we investigated whether the widely accepted hypothesis that specialist exhibit higher *k*_cat_ values compared to generalist enzymes holds true in Yeast-MetaTwin^27, 47^. All three prediction methods confirmed that Yeast-MetaTwin aligns with this perspective (Supplementary Figure 11).

Furthermore, we explored the underground reaction proportion among different reaction types. We counted the reaction numbers involving different EC numbers in Yeast8 (known metabolism) and Yeast-MetaTwin (known and underground metabolism). We found that the EC5.X.X.X (isomerase) category has the lowest proportion of explored metabolism, with 4.3% explored. In contrast, the explored proportion is highest for EC6.X.X.X (ligase), at 43.9%. These distributions are also consistent when considering the yeast annotation in UniProt as the known metabolism (Supplementary Figure 12). A previous study has shown that isomerase (EC5.X.X.X) binds the least tightly to its substrate, whereas transferase (EC2.X.X.X) and ligase (EC6.X.X.X) bind the most tightly to their substrates, which is in good agreement with our results and indicates that the tightness of substrate-enzyme binding affects enzyme promiscuity^48^.

### Yeast-MetaTwin predicts by-products for native chemicals produced in yeast

In cell factories, by-products may be produced alongside the desired product, representing wasteful use of resources, impeding efficiency and yield in the production of the desired product^49^. As cell factories often have altered metabolism due to genetic and metabolic engineering, predicting by-products is challenging, especially those related to product degradation by potential promiscuous and/or uncharacterized enzymes (Fig. 4a). Here, with the Yeast-MetaTwin model, which comprehensively describes enzyme promiscuity and metabolite conversions, we can locate potential product degradation reactions in yeast. For previously published 48 native products produced in yeast^50^, we counted reactions in which those products can serve as reactant in both Yeast8 and Yeast-MetaTwin and identified the potential degradation by-products for those native products. Among all tested native products, amino acid-derived products and specific organic acids, such as pyruvate and 2-oxoglutarate, exhibited the highest number of degradation reactions (Fig. 4b). This can be explained by the fact that those products are central metabolites, which usually carry large fluxes and could therefore, have more tendency to activate its enzyme’s promiscuities in the long history of evolution. These products were also predicted to be hub metabolites as discussed in the previous sections. Furthermore, we observed consistency between the predicted by-products in Yeast-MetaTwin and experimental reports (Fig. 4b). For instance, the conversion of L-glutamate to L-glutamate 5-semialdehyde catalyzed by Put2 (YHR037W)^51^, which is part of the proline metabolism pathway, was absent in Yeast8 but captured in Yeast-MetaTwin. Additionally, there is a recent report of the decarboxylation of lysine to produce cadaverine in yeast as a non-specific substrate activity of ornithine decarboxylase (Spe1, YKL184W)^52^. In Yeast-MetaTwin, this reaction was also successfully predicted as the by-product production reaction for lysine production and the gene YKL184W was also successfully identified as the encoding gene. Moreover, homocysteine S-methyltransferase (Sam4, YPL273W) is involved in the conversion of homocysteine and S-adenosylmethionine to methionine, thereby regulating the methionine/S-adenosylmethionine ratio^53^. This reaction, along with its catalyzing enzyme Sam4, has been successfully captured in Yeast-MetaTwin, it is considered a potential by-product formation reaction during methionine production (Fig. 4b). Furthermore, the transaminase function of Aro9 (YHR137W)^54^, which catalyzes the reaction from 2-oxoglutarate and L-kynurenine to produce kynurenate and L-glutamate, was also successfully captured in Yeast-MetaTwin but not in Yeast8.

**Figure 4.**
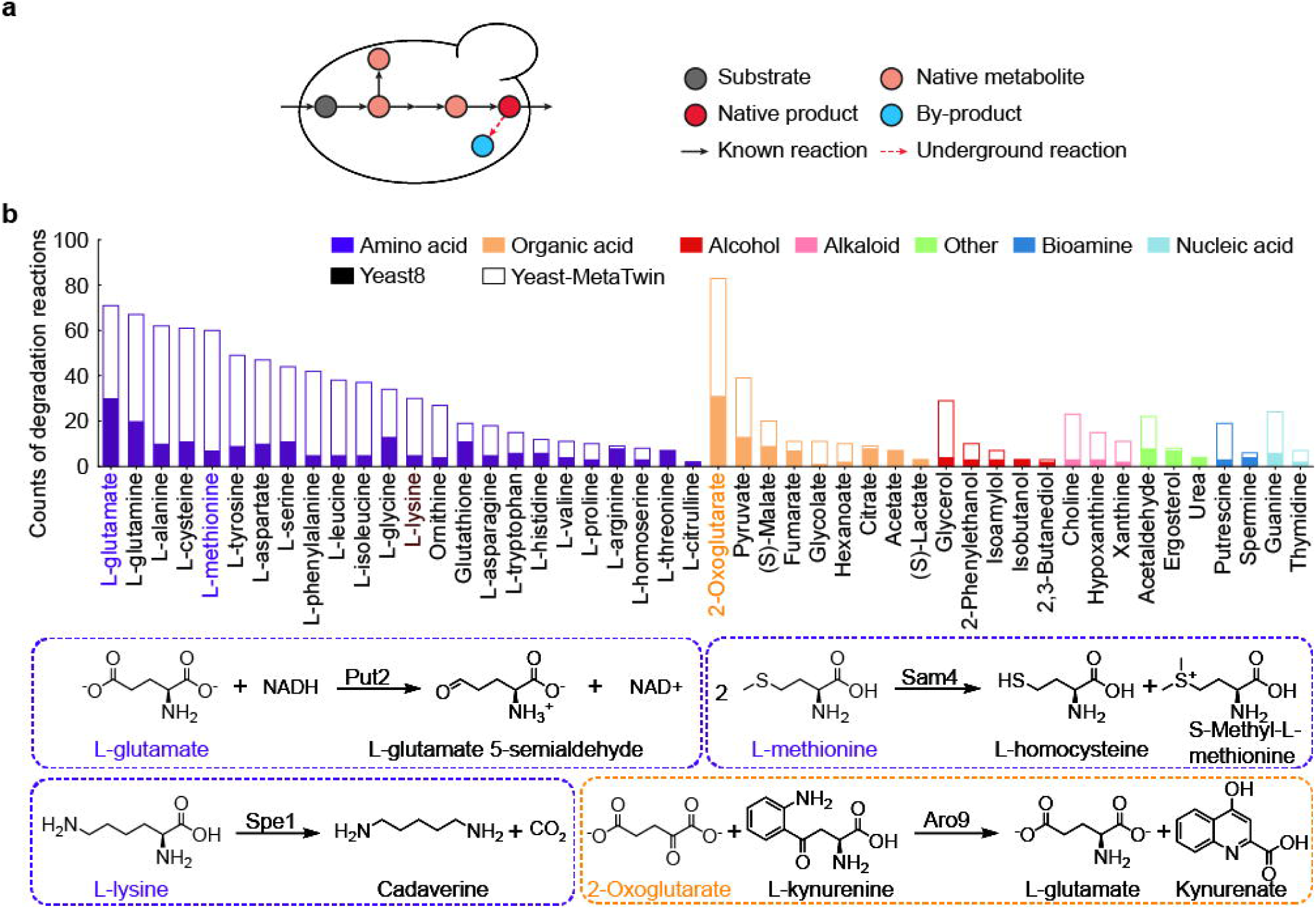
Predicted By-products degradation reactions of native products in *S. cerevisiae*. (a) The scheme for native product production involves underground degradation reactions. (b) Degradation reactions for 48 native products in *S. cerevisiae* were predicted using Yeast8 and Yeast-MetaTwin, as well as literature-reported cases specific to Yeast-MetaTwin. Put2, delta-1-pyrroline-5-carboxylate dehydrogenase; Sam4, homocysteine S-methyltransferase 2; Spe1, ornithine decarboxylase; Aro9, aromatic amino acid aminotransferase 2.

### By-product identification for heterologous chemical production in yeast

By-product formation is an even more common issue in production of heterologous products and remains to be systematically explored (Fig. 5a). We adapted our Yeast-MetaTwin pipeline to predict the potential degradation reactions for 44 heterologous products collected from previous studies^50^. We predicted an average of 12 conversion reactions for each product, and found that terpenoids, aromatic compounds, and flavonoids have a high potential to be degraded in yeast (Fig. 5b). This suggests that in terms of metabolic stability and yield, these compound classes require more careful optimization and engineering to minimize degradation and enhance their production efficiency in yeast. As for all by-product conversions, we not only identified reactions but also gave candidate genes corresponding to those degradation reactions. We also observed consistencies in experimental reports with our predictions (Fig. 5b). Notably, our methods predicted the two by-products conversions of geranial^55^ and citronellol^56^ in geraniol production and annotated the latter conversion to be associated with an enzyme encoded by Oye2 (YHR179W) as has been experimentally reported^56^.Conversions of limonene to carveol^55^ and psilocybin to psilocybin^57^ were also successfully predicted. Furthermore, L-phenylacetylcarbinol (L-PAC) was produced from acetaldehyde (ACA) with the extra addition of benzaldehyde (BZA)^58^. During this process, three main by-products were identified: 1-phenyl-1,2-propanediol (PAC-diol), which is degraded from L-PAC; benzoic acid (BZOA), resulting from BZA oxidation; and benzyl alcohol (BZOH), formed via BZA reduction^58^. Our methods successfully predicted these three by-products in L-PAC production, along with the corresponding gene annotations (Fig. 5b).

**Figure 5.**
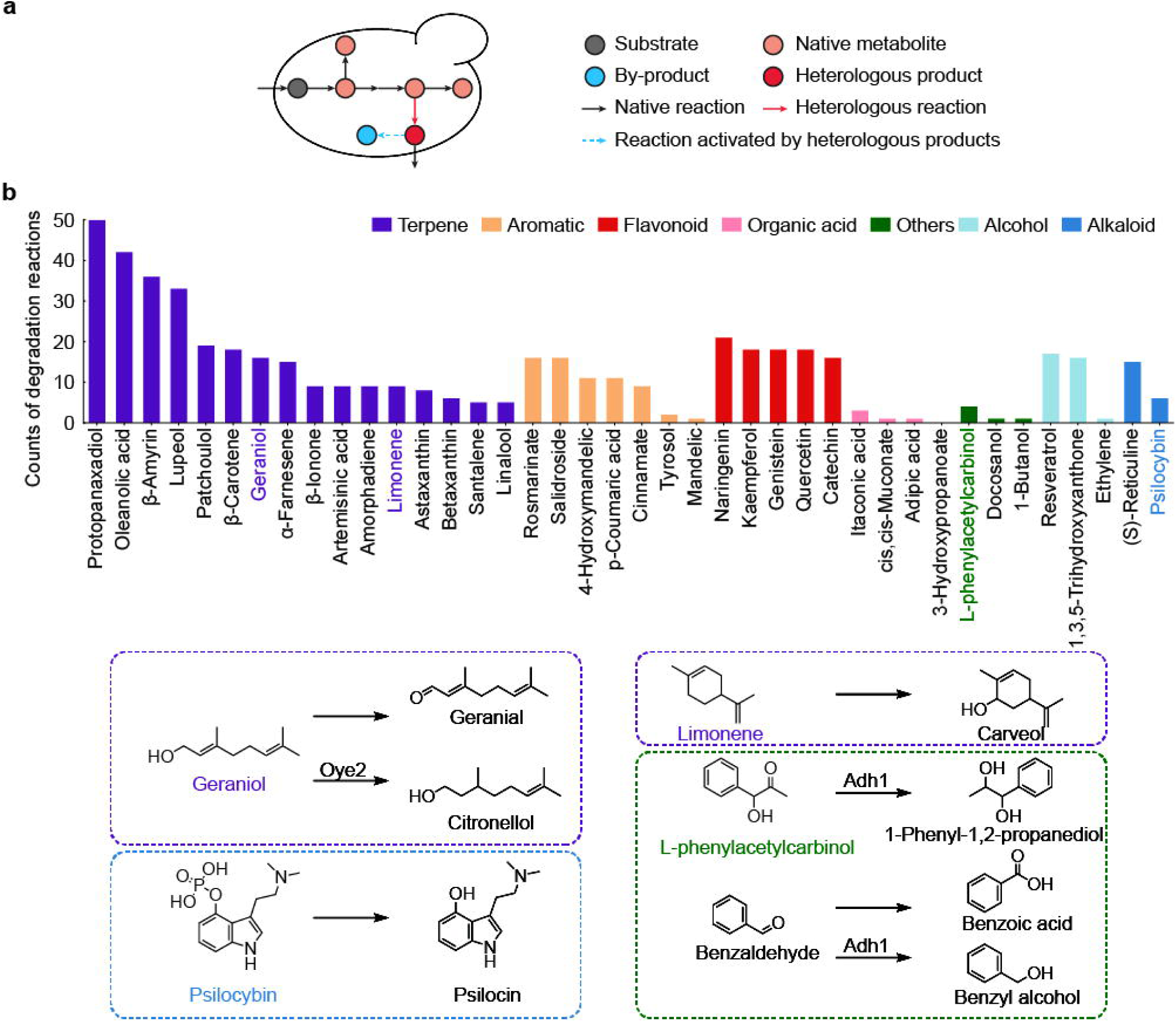
Predicted By-products degradation reactions of heterologous products in *S. cerevisiae*. (a) The scheme for heterologous product production involves underground degradation reactions. (b) Degradation reactions for 44 heterologous products in *S. cerevisiae* were predicted using Yeast8 and Yeast-MetaTwin, as well as literature-reported cases specific to Yeast-MetaTwin. Oye2, NADPH dehydrogenase 2. ADH1, Alcohol dehydrogenase 1.

## Discussion

In this study, we developed a workflow to reconstruct comprehensive metabolic networks, which utilizes metabolome data to constrain the reaction expansion process, ensuring that all metabolites involved in novel reactions are found in the metabolome. This approach streamlines the reaction expansion to a single step, effectively addressing the combinatorial explosion that has previously been encountered in the retrobiosynthesis pipelines^59^. Moreover, we utilized advanced deep learning enzyme annotation methods to further trim the reaction pool^31, 32^, removing reactions that cannot be annotated with enzymes. This ensures effective establishment of a comprehensive underground metabolic network. Our effort resulted in the development of Yeast-MetaTwin, the comprehensive metabolic twin model of *S. cerevisiae*. Yeast-MetaTwin exhibited a significant enhancement in the coverage of gene, metabolite and reactions, encompassing 84% of predicted metabolic enzymes and 92% of the yeast metabolome, which improves the predictive performance of growth phenotypes and gene essentiality. The comprehensive metabolic network enhances metabolic factory redesign, by identifying traditionally unpredictable by-product pathways. We investigated the kinetic differences between the known network (Yeast8) and underground network (predicted new reactions in Yeast-MetaTwin) using deep learning methods. Our study implied that underground metabolism is dominated by variations in *K*_m_ and not *k*_cat_ values, which confirms previous assumptions based on sparse experiment measurements^60^.

Ever since the publication of the first GEM and the standard protocol of GEM reconstruction^61, 62^, regardless on top-down or bottom-up, manual or automatic^63–66^ approach, GEM reconstructions have always been limited by the known biochemical reactions and characterized metabolic enzymes. Moreover, the model reconstruction process is frequently required to (at least partially) rerun to keep up with new knowledge, which requires extensive effort^67^. In contrast, the approach presented here brings a new promise to break this boundary to reconstruct the GEM with the reaction mechanism and the deep learning enzyme annotation methods.

Moreover, our pipeline can be directly extended to other organisms, such as *Escherichia coli*^68^ and *Homo sapiens*^69^, with the corresponding curated metabolome databases ECMDB (*Escherichia coli* Metabolome Database)^70^ and HMDB (Human Metabolome Database)^71^, which can advance the drug discovery and human health engineering. Additionally, it can be applied to microbial consortia with metabolome data, such as human gut microbiota, which can be used to study complex interactions within microbial communities and their impact on host metabolism^72–74^. With the advancement of future high-throughput metabolome techniques^75–77^, it also brings the promise to reconstruct the comprehensive metabolic twin model for all species.

Even though the resulted Yeast-MetaTwin presents superior metabolic coverage and performance, it faces limitations. For example, we were unable to find enzyme-annotated reactions to connect 267 metabolites (1.7% of the yeast metabolome) to the Yeast-MetaTwin model without introducing non-yeast-metabolome metabolites. This could be due to the limitations in our current knowledge of biochemical reactions, indicating that there exist novel reaction mechanisms which are not included in the current reaction rule pool. Deep learning-based retrosynthesis methods is able to partially solve this issue^23^, but difficulties arise since those methods usually predict one to one mapping of reactants to products, without considering the stoichiometry and element balance required for the GEM reconstruction and flux simulation. Alternatively, errors in metabolome data could also be a factor, such as incomplete data or false positives arising from non-enzymatic reactions during sample handling or interactions with solvents and plastics^78, 79^.

Another limitation encountered in our research is the enzyme annotation method. Despite the boom of enzyme annotation methods^31, 32, 80^, particularly with advancements in deep learning, current methods still have substantial room for improvement. For instance, existing enzyme annotation approaches typically link proteins to GO terms^81, 82^, EC numbers^31, 32, 80^, substrates^33^, or use kinetic parameters like *k*_cat_ and *K*_m_ to characterize enzyme functions^27–29^. However, there remains a gap in directly annotating enzymes to new reactions. Although some methods attempted to directly annotate enzymes to reactions by encoding reaction formulas as embeddings, they could not be generalized to novel reactions and thus could not identify enzymes for novel reactions^29^. Moving forward, the development of methods that can directly annotate proteins to novel reactions holds promise for enhancing our pipeline’s capabilities^59, 83^.

In summary, we developed a comprehensive metabolic model reconstruction pipeline integrating retrobiosynthesis and deep learning enzyme annotation methods, along with a delicate design to avoid combinatorial explosion issues. This pipeline resulted in the reconstruction of Yeast-MetaTwin, a comprehensive metabolic twin model for yeast that encompasses both known and underground metabolism. Notably, Yeast-MetaTwin showed significant improvements in metabolite coverage, gene-associated reactions, and phenotype prediction. Furthermore, our study identified affinity, rather than turnover number, as a key factor distinguishing known and underground metabolism, shedding light on understanding yeast’s metabolic potential and adaptive responses in diverse environmental conditions. Last, we validated the model’s predictive power in cell factory design, demonstrating its capability in by-product prediction and enhancing cell factory efficiency. Understanding underground metabolism contributes significantly to our knowledge of biological metabolism, facilitates the integration of omics data, and advances metabolic engineering for cell factories.

## Methods

### Dataset preparation for yeast metabolite pool

First, we obtained metabolite entries with SMILES information from YMDB (Yeast Metabolome Database, https://www.ymdb.ca/)^12^. These metabolites were then classified using the ClassyFire^38^, which utilizes chemical structures and structural features to categorize compounds. Simultaneously, for metabolites in the Yeast8.7.0^13^, we used APIs to retrieve SMILES information from various databases based on the CHEBI ID^84^, KEGG ID^85^, and MetaNetX ID^39^ annotated in the model. Finally, metabolites from the YMDB database were merged with Yeast8 metabolites to create a yeast metabolite pool, using SMILES mapping.

### Reaction rule extraction

We extracted all reactions in the MetaNetX database^39^, excluding transport and unbalanced reactions, and removed H^+^ and the pairs of metabolites involved in each reaction (e.g., atp --> adp + p_i_). Subsequently, we used RXNMapper for atomic mapping^86^ of these reactions and employed RDChiral for rule extraction^87^. The atomic radius for rule extraction was set to 1, except for reactions involving bond formation or breaking within a single molecule. In such cases, the radius was adjusted to the minimum radius that ensures that the substructures extracted from the rules matched the number of metabolites in the template reaction.

### Reaction predictions based on reaction rules and metabolite pool

The extracted rules were then applied to the yeast metabolite pool to predict reactions using RunReactants function in RDKit (https://www.rdkit.org/). All predicted novel reactions inherited the first three EC number digits (EC X.X.X.-) from their corresponding template reactions^40, 41^. The criteria for reaction expansion were as follows: the metabolite must contain the substructure specified by the reaction rule, and the top 50 metabolites with a structure similarity greater than 0.3 compared to the metabolites in the template reaction were selected. For rules that include co-substrates, the metabolites meeting the criteria were combined to generate a set of reactions. Those parameters were optimized through sampling (Supplementary Figure 3). Reaction formulas were then modified to add the removed currency metabolites and the metabolite pairs to meet stoichiometry and element balance. The detailed processes of applying rules to predict reactions is illustrated in Supplementary Figure 1. We applied the rule in the forward direction and reverse direction to fully cover the synthesis and degradation of all metabolites in the pool. Due to the imbalanced distribution of lipid and non-lipid metabolites in YMDB, we separated the lipids and non-lipids in the reaction predictions process. As for the large amount lipid-related metabolites, all metabolites with similarity greater than 0.3 were selected for combination, rather than just the top 50, to ensure sufficient retrobiosynthesis coverage for lipid metabolites.

### Reaction network filtering

The retrobiosynthesis process generated numerous reactions, all of which included yeast metabolites either as substrates or products. To ensure the integrity and connectivity of these reactions within the Yeast8 framework, a network filtering process was implemented. This process includes two main aspects: 1) eliminating reactions that contain metabolites not in the yeast metabolite pool and 2) ensuring that reactions successfully connect to Yeast8. The first aspect removes large proportions of reactions that contain non-yeast-metabolome metabolites, limiting to a reasonable reaction space for yeast. As for the second aspect to filter out the underground network that can be successfully connected with Yeast8, a four-step network filtering process was devised based on the structure of the graph:

a. Firstly, all metabolites in the predicted reactions were compared to Yeast8 metabolites. A score of 1 was assigned to all metabolites already present in Yeast8.
b. When traversing each reaction, if all child nodes (reactants) of a reaction reached a score of 1, it indicated that the reaction is successfully connected to Yeast8. Therefore, the score of the root node (products) was updated to 1 and broadcasted to all reactions.
c. Step b was repeated until no new metabolite scores were updated to 1.
d. Reactions were filtered out where all metabolite scores were equal to 1, resulting in a yeast connected network linking from the Yeast8 to the underground network.

### Reaction annotation

To annotate the reaction pool generated by retrobiosynthesis and integrate it into the Yeast8, we employed deep learning methods^31^ to predict EC numbers for the entire yeast genome and matched these predictions with the EC numbers of the reaction pool, thereby annotate those reactions with enzymes. For multiple genes predicted to be associated with the same EC number, we applied a logical "OR" relationship when integrating them into the GEM. We evaluated the accuracy of two EC number prediction methods, CLEAN^31^ and DeepECtransformer^32^, and found that CLEAN had higher recovery but also a higher false discovery rate, while DeepECtransformer had lower recovery but a lower false discovery rate (Supplementary Figure 4a-b). To balance accuracy and false discovery rate, we ultimately combined both methods: using CLEAN’s prediction results, along with DeepECtransformer to determine if the protein is a metabolic enzyme thus maintaining high accuracy while reducing the false discovery rate.

In particular, this step filtered out two main types of reactions that cannot find enzyme associations. Firstly, no enzyme in yeast is annotated to the EC number of the reaction. Secondly, the reaction does not have an EC number, making it impossible to assign enzymes. This situation is caused by the missing EC number in template reactions in the MetaNetX reaction database. If these reactions are permitted, four metabolites can be incorporated into Yeast-MetaTwin via four reactions that lack enzyme annotations (Supplementary Table 3).

### Prediction of dynamic parameters

To evaluate the differences in kinetic parameters between underground and known network, we used two *k*_cat_ prediction methods DLKcat^27^and UniKP^28^ to predict the *k*_cat_ values of enzyme-substrate pairs, and used Boost_KM^30^ and UniKP^28^ to predict the *K*_m_ of enzyme-substrate pairs. These methods use the SMILES of individual reactants and enzyme sequences as input. Moreover, we used TurNuP to predict *k*_cat_ values^29^. Unlike methods based on enzyme-substrate pairs, TurNuP utilizes the complete SMILES of the reaction along with enzyme sequences as input^29^.

### Gene essentiality simulation

Gene essentiality was simulated by blocking the reactions associated with individual genes and setting growth as the objective function. A rich medium was set to match the experimental conditions, and a growth cutoff of less than 0.1*max biomass was used to determine essentiality. Accuracy and the confusion matrix were calculated. The essential gene list was obtained from the Yeast Deletion Project (available at http://www-sequence.stanford.edu/group/yeast_deletion_project/downloads.html), which was generated from experiments using a complete medium^13^.

### Amino acids biosynthesis robustness simulation

The amino acid synthesis pathways and their corresponding synthetic genes were identified using pFBA^88^ simulations, with specific amino acid synthesis set as the objective function under minimal medium conditions. Identified genes which can carry flux except those in glycolysis are determined as synthetic genes for each amino acid. Then, for each amino acid, a randomly chosen set of genes from its synthesis pathways was in silico knocked out by blocking associated reactions.

The specific amino acid synthesis was then reassessed as the objective function. This process was repeated 100 times for each case to evaluate the impact. A yield cutoff of less than 0.1* max biomass was employed to determine the loss of amino acid synthesis ability.

### Prediction of byproduct for heterologous products

The method for predicting degradation pathways of heterologous products is similar to predicting underground metabolic reactions. It incorporates both yeast metabolites and target heterologous products into the metabolite pool for retrobiosynthesis, then identifies reactions with the target exogenous product as a reactant. Then the enzyme annotation pipeline was used to annotate possible genes for the degradation.

### Data availability

MetaNetX database (https://www.metanetx.org/) was used in the retrobiosynthesis reaction rules extraction. UniProt database (https://www.uniprot.org/) was used to evaluate the workflow performance. RetroPathRL (https://github.com/brsynth/RetroPathRL) and ASKCOS (https://github.com/itai-levin/chemoenzymatic-askcos) were used to illustrate metabolites that not be well-suited for template-based retrobiosynthesis methods. Methods including CLEAN (https://github.com/tttianhao/CLEAN), DeepECtransformer (https://github.com/kaistsystemsbiology/DeepProZyme) were used in EC number prediction. Methods including DLKcat (https://github.com/SysBioChalmers/DLKcat), UniKP (https://github.com/Luo-SynBioLab/UniKP), TurNuP (https://github.com/AlexanderKroll/kcat_prediction) were used in *k*_cat_ prediction. Methods including UniKP (https://github.com/Luo-SynBioLab/UniKP) and Boost_KM (https://github.com/AlexanderKroll/KM_prediction) were used in *K*_m_ prediction. The authors declare that all data supporting the findings and for reproducing all figures of this study are available within the paper and its Supplementary Information. Source data are provided with this paper. All data used in this study can be found in https://github.com/LiLabTsinghua/Yeast-MetaTwin.

### Code availability

To facilitate further use, we have made all the codes and detailed instructions available in our GitHub repository, located at https://github.com/LiLabTsinghua/Yeast-MetaTwin.

## Acknowledgements

This study was financially supported by the National Key R&D Program of China (2023YFA0913900), the Shenzhen Medical Research Fund (A2303026) and the Shenzhen Science and Technology Program (KJZD20230923114415032).

## Author Contributions

F.L., J.N., Y.C. and K.W. designed the research. K.W., H.L. and M.S. performed the research. K.W., H.L., F.L., Y.C., E. J. K. and J.N. analyzed the data. F.L. and K.W. wrote the paper. K.W., R. M., and Y. J. performed by-product analysis. All authors approved the final paper.

