## Supplementary Figure for "Yeast-MetaTwin for Systematically Exploring Yeast Metabolism through Retrobiosynthesis and Deep Learning"

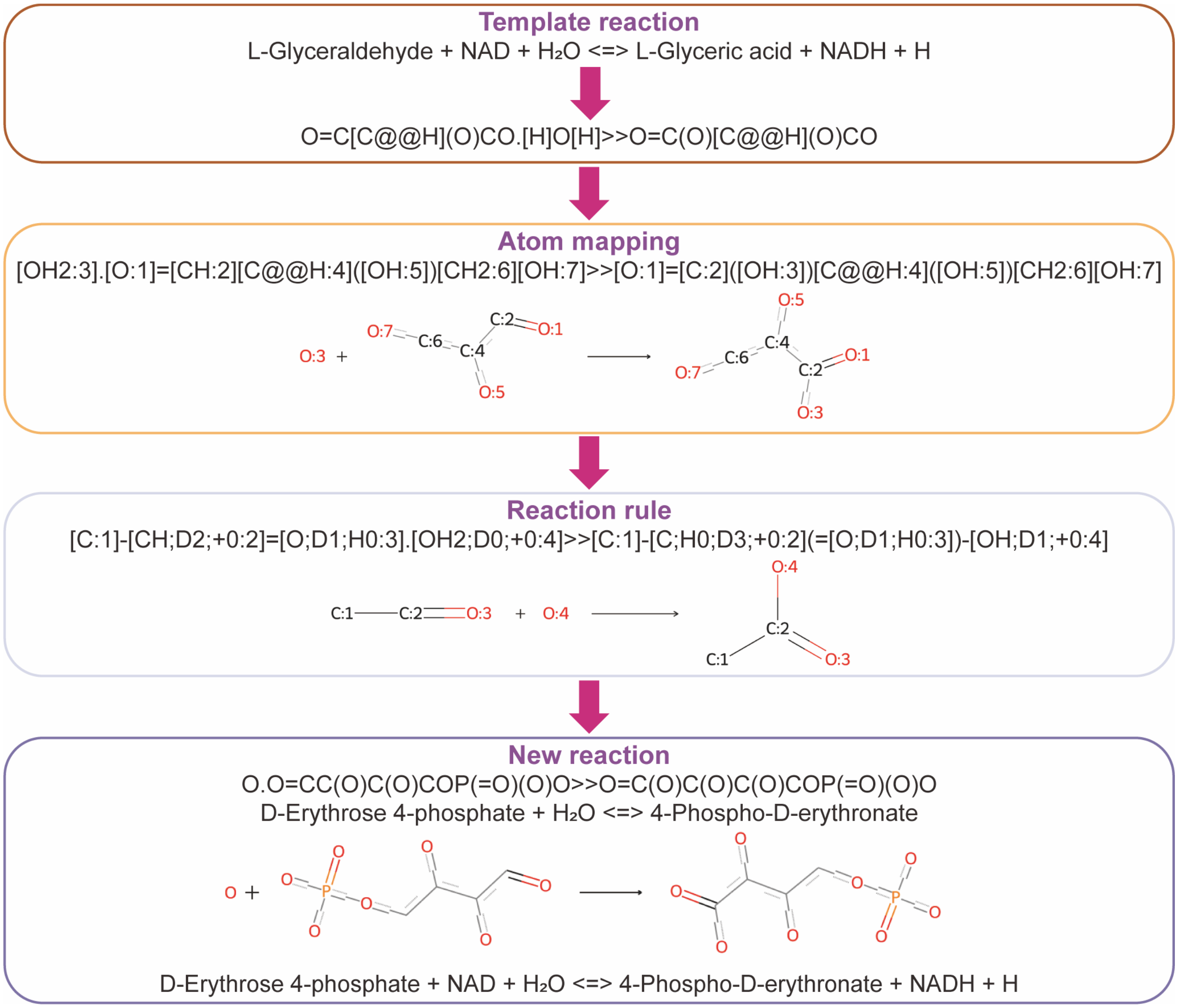


**Supplementary Figure 1** Schematic representation of reaction rule extraction and reaction prediction.

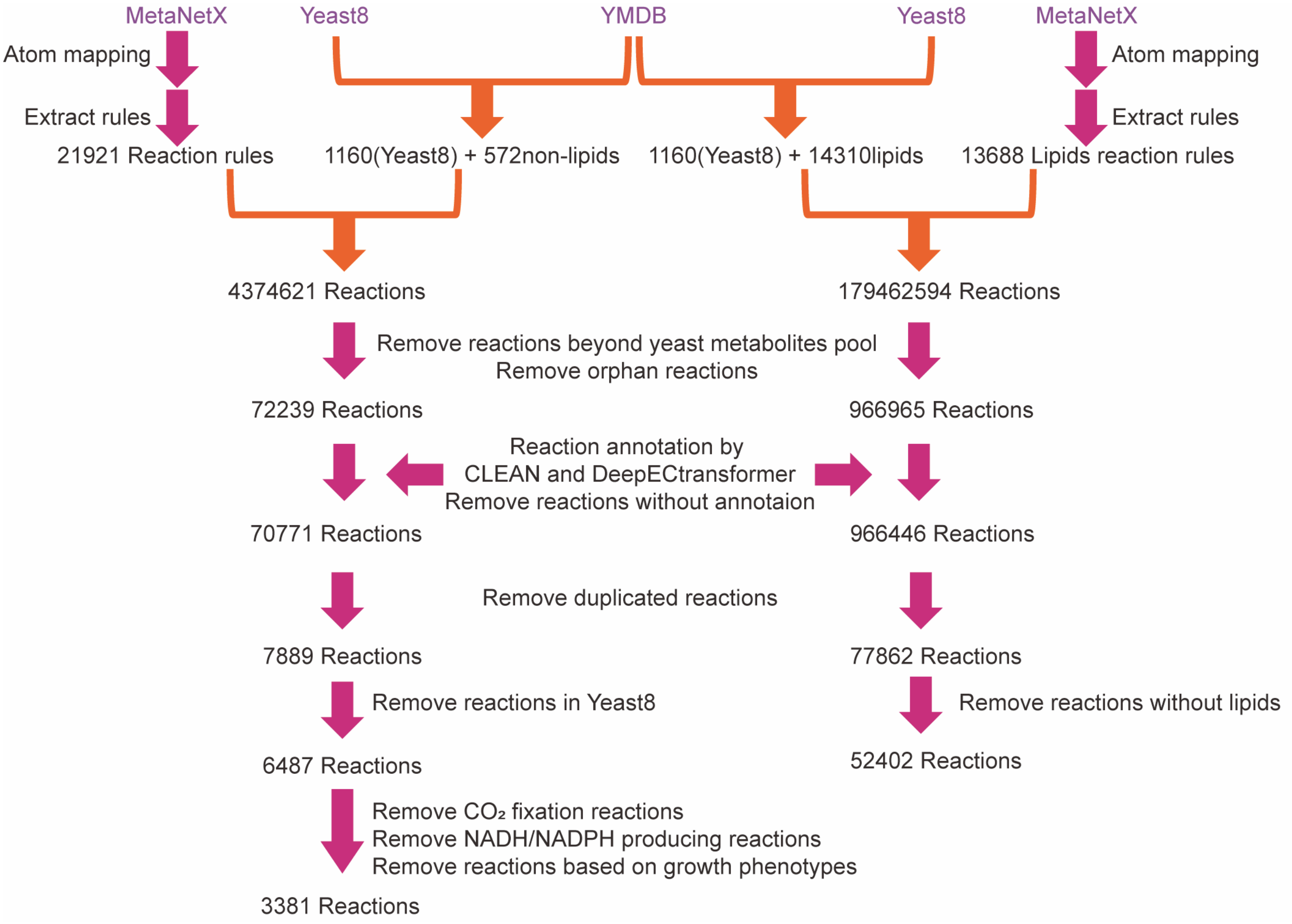


**Supplementary Figure 2** Detailed processes for reaction prediction and annotation.

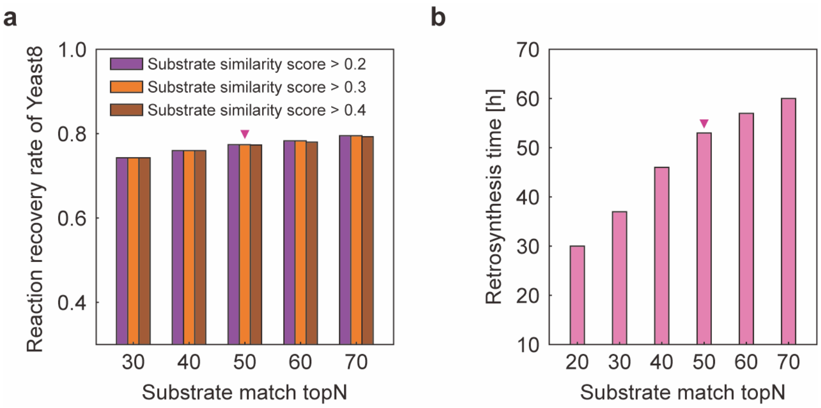


**Supplementary Figure 3** Performance and runtime under different substrate similarity cutoffs for the retrosynthesis. (a) Reaction recovery rate of Yeast8 using different substrate similarity cutoffs. (b) Runtime under different substrate similarity cutoffs. Substrate similarity cutoffs refer to the counts of metabolites that are most similar to the substructures within the reaction rules when performing reaction prediction.

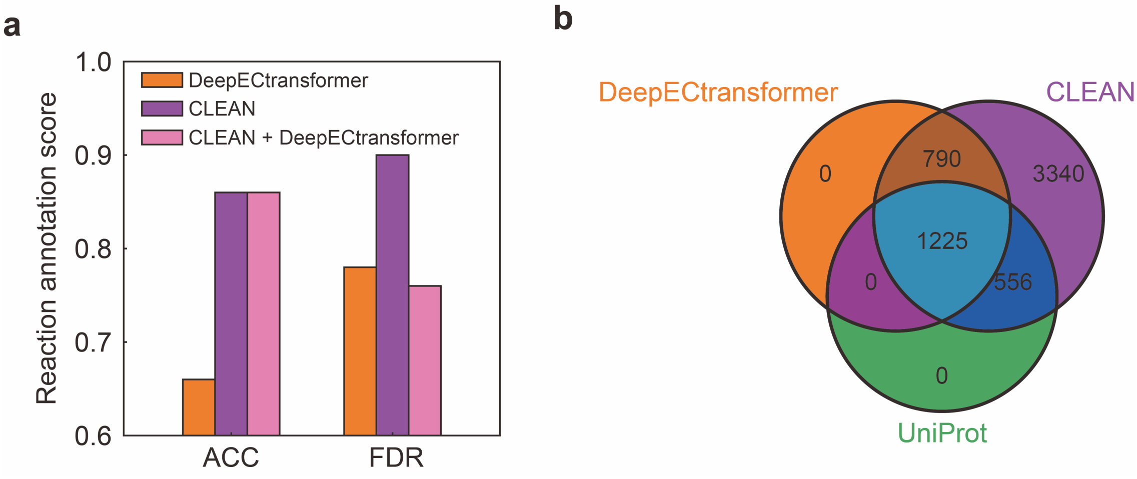


**Supplementary Figure 4** Performance of reaction annotation using different deep learning methods for EC number prediction. (a) The reaction annotation performance of CLEAN[1], DeepECtransformer[2], and their combination. The combination of CLEAN with DeepECtransformer is achieved by excluding non-enzyme genes predicted by DeepECtransformer from the results of CLEAN. (b) Comparison of enzyme annotation sets using CLEAN, DeepECtransformer, and UniProt database. ACC: the recovery rate of correctly predicted Yeast8 reactions that have consistent enzyme annotations identified within Yeast8. FDR: one minus the ratio of Yeast8 enzymes among all predicted enzymes for correctly recovered reactions. This metric indicates the number of isozymes identified per reaction.

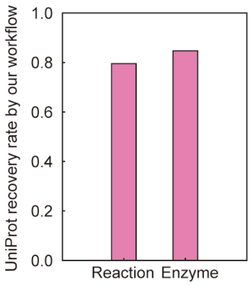


**Supplementary Figure 5** The reaction recovery rate and reaction annotation accuracy in UniProt by our process. Reaction stands for the reaction recovery rate, while the enzyme stands for the successful reaction annotation rate.

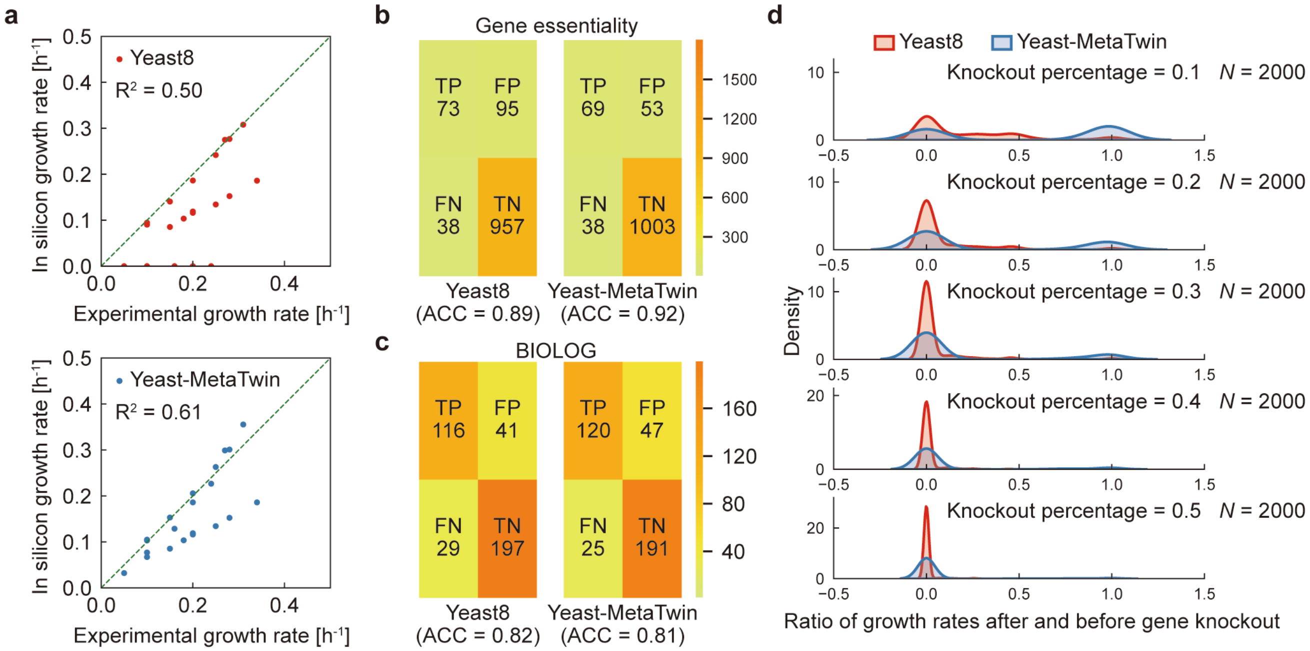


**Supplementary Figure 6** Model performance comparison of Yeast8 and Yeast-MetaTwin. (a) Growth prediction in Yeast8 and Yeast-MetaTwin, with each point representing a specific growth condition. Experimental data are from previous work[3]. (b) Confusion matrix for gene essentiality prediction in Yeast8 and Yeast-MetaTwin. Experimental data are from previous work[4]. (c) Confusion matrix for BIOLOG substrates utilization prediction in Yeast8 and Yeast-MetaTwin. Experimental data are from previous work[3]. TP: true positive, TF: true negative, FP: false positive and FN: false negative. (d) The amino acid synthesis capacity retained by Yeast8 and Yeast-MetaTwin under various gene knockout percentage. The gene knockout percentage represent the percent of gene knockouts across 20 different amino acid synthesis pathways, with each amino acid being sampled 100 times.

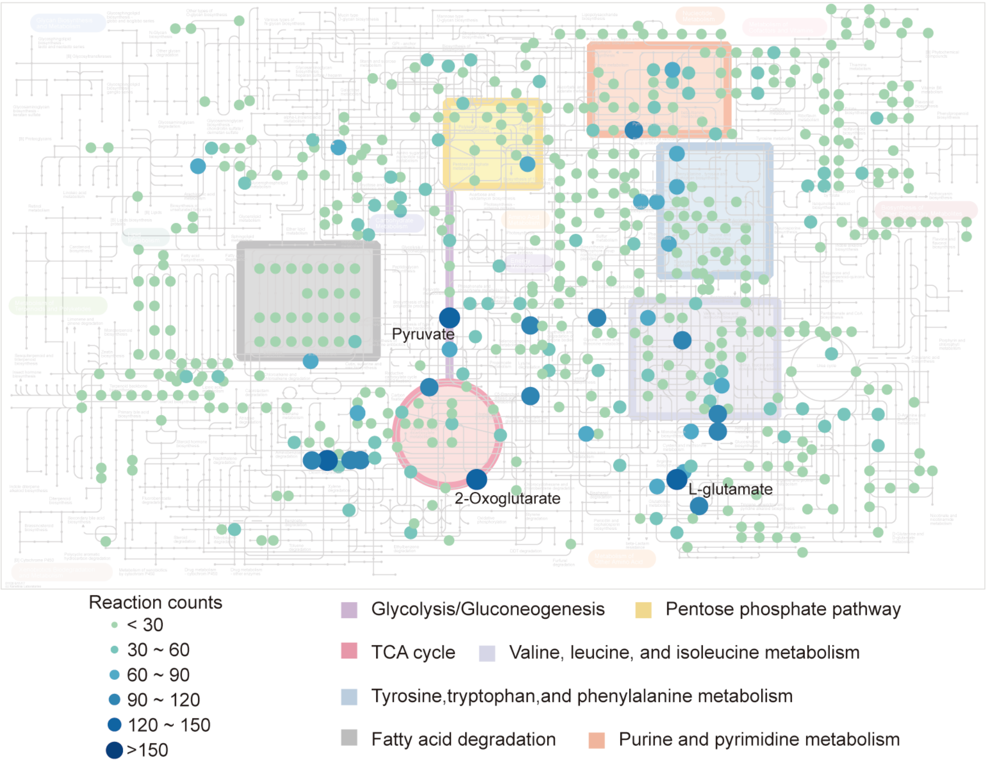


**Supplementary Figure 7** Reaction counts for each metabolite in Yeast-MetaTwin (without lipids). In the metabolic maps, nodes represent metabolites from Yeast8 involved in underground metabolism, with their color and size corresponding to the number of reactions they participate in.

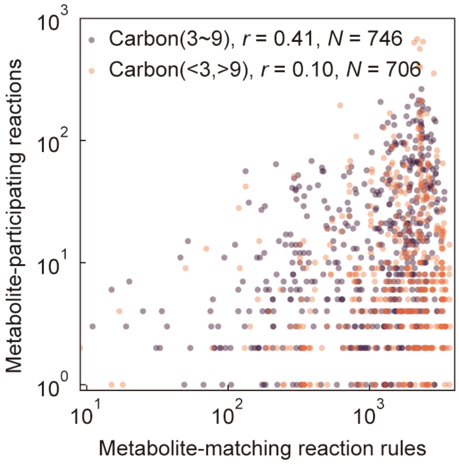


**Supplementary Figure 8** Correlation between the number of metabolite-matching reaction rules and the number of participating reactions in Yeast-MetaTwin.

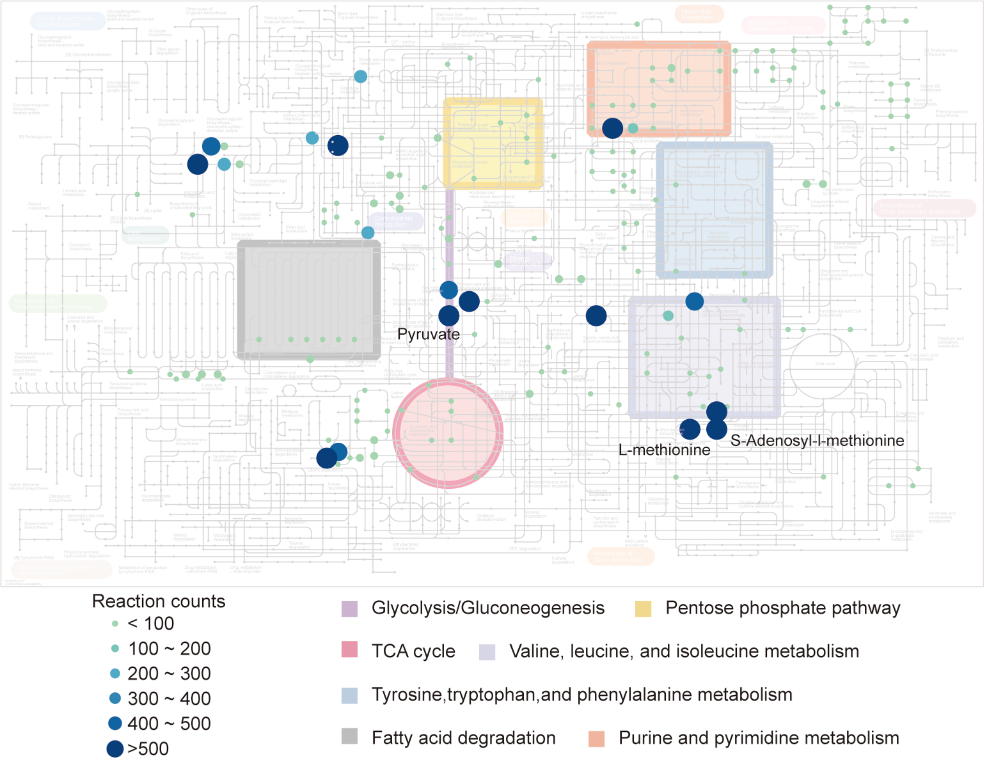


**Supplementary Figure 9** Reaction counts for each metabolite in Yeast-MetaTwin (only lipids). In the metabolic maps, nodes represent metabolites from Yeast8 involved in underground metabolism, with their color and size corresponding to the number of reactions they participate in.

85

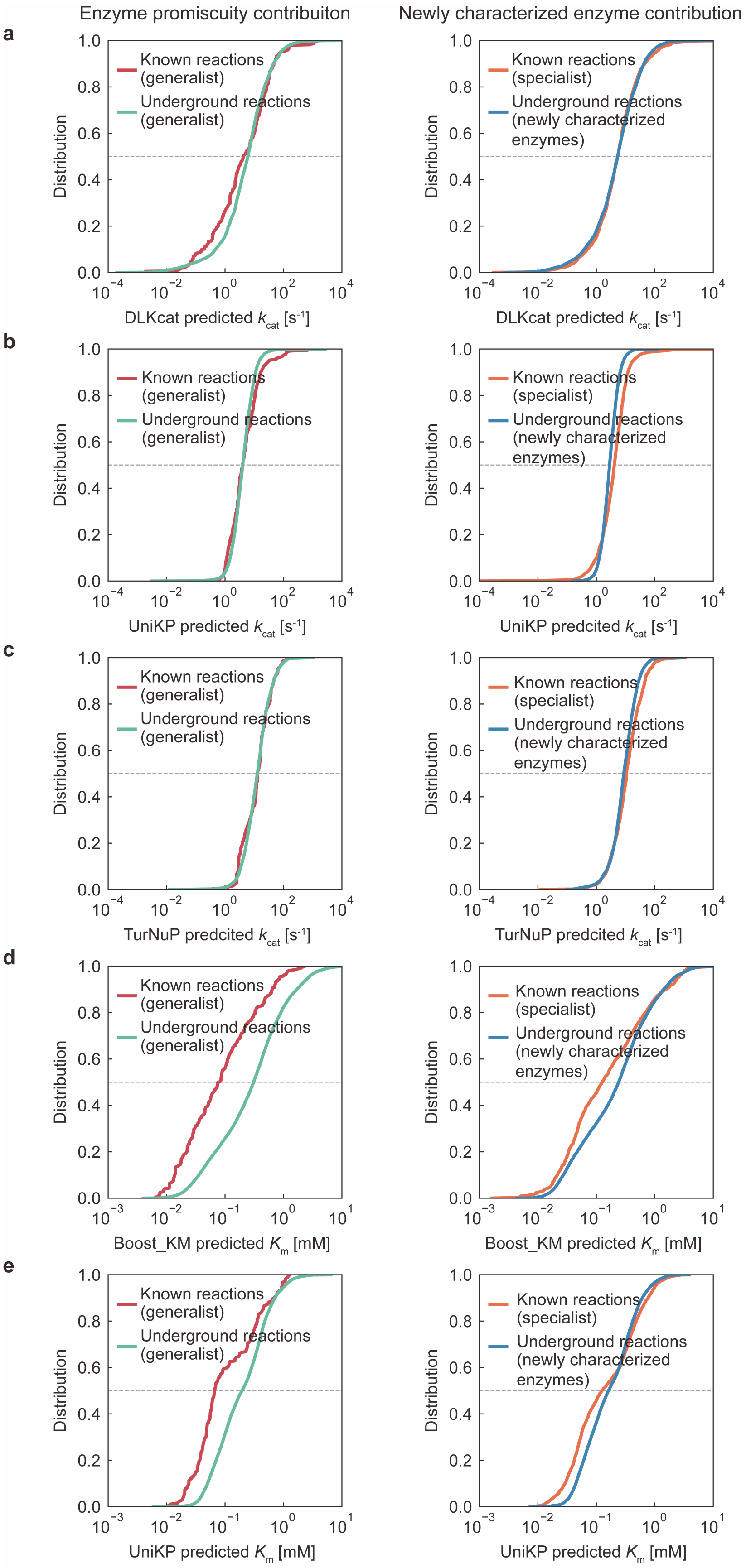
Enzyme promiscuity contribuiton
Newly characterized enzyme contribution
a
b
c
d
e
Known reactions (generalist)
Underground reactions (generalist)
Known reactions (specialist)
Underground reactions (newly characterized enzymes)
Distribution
DLKcat predicted $k_{cat}$ [s$^{-1}$]
UniKP predicted $k_{cat}$ [s$^{-1}$]
UniKP predcited $k_{cat}$ [s$^{-1}$]
TurNuP predcited $k_{cat}$ [s$^{-1}$]
Boost_KM predicted $K_m$ [mM]
UniKP predicted $K_m$ [mM]

**Supplementary Figure 10** Kinetic parameter prediction for the generalist enzymes and the newly characterized enzymes. (a) distribution of $k_{cat}$ values predicted by DLKcat[5], (b) distribution of $k_{cat}$ values predicted by UniKP-$k_{cat}$[6], (c) distribution of $k_{cat}$ values predicted by TurNuP[7], (d) distribution of $K_m$ values predicted by Boost_KM[8], and (e) distribution of $K_m$ values predicted by UniKP-$K_m$[6]. The left side compares the kinetics of promiscuous enzymes (generalists in Fig. 3d) in underground reactions versus previously known Yeast8 reactions, highlighting the impact of enzyme promiscuity. The right side compares the kinetics of newly characterized enzymes with specialist enzymes in Yeast8, illustrating the impact of these newly characterized enzymes.

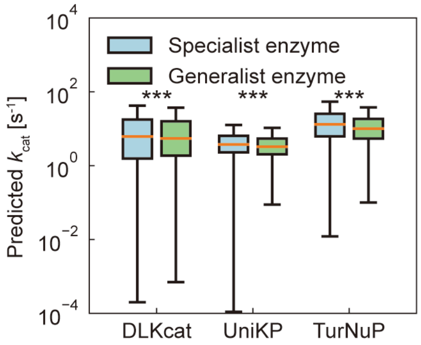


**Supplementary Figure 11** Predicted $k_{\text{cat}}$ values for reactions catalyzed by generalist and specialist enzymes in the Yeast-MetaTwin. The two-sided Wilcoxon rank sum test was used to calculate *P* value. *** means *P* value < 0.001 in the correlation test analysis.

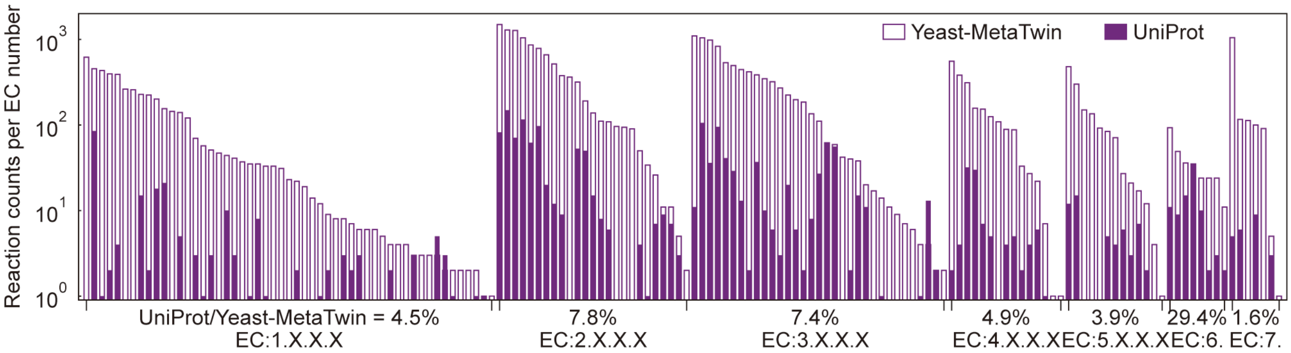


**Supplementary Figure 12** Proportions of explored reactions in different EC number categories in UniProt database annotation and Yeast-MetaTwin (without lipids).
